# Repeated evolution of inactive pseudonucleases in a fungal branch of the Dis3/RNase II family of nucleases

**DOI:** 10.1101/2020.07.30.229070

**Authors:** Elizabeth R. Ballou, Atlanta G. Cook, Edward W.J. Wallace

**Affiliations:** Institute for Microbiology and Infection, School of Biosciences, University of Birmingham; Wellcome Centre for Cell Biology, School of Biological Sciences, University of Edinburgh; Institute for Cell Biology, School of Biological Sciences, University of Edinburgh

## Abstract

The RNase II family of 3′-5′ exoribonucleases are present in all domains of life, and eukaryotic family members Dis3 and Dis3L2 play essential roles in RNA degradation. Ascomycete yeasts contain both Dis3 and inactive RNase II-like “pseudonucleases”. These function as RNA-binding proteins that affect cell growth, cytokinesis, and fungal pathogenicity. Here, we show how these pseudonuclease homologs, including *Saccharomyces cerevisiae* Ssd1, are descended from active Dis3L2 enzymes. During fungal evolution, active site mutations in Dis3L2 homologs have arisen at least four times, in some cases following gene duplication. The N-terminal cold-shock domains and regulatory features are conserved across diverse dikarya and mucoromycota, suggesting that the non-nuclease function require this region. In the basidiomycete pathogenic yeast *Cryptococcus neoformans*, the single Ssd1/Dis3L2 homolog is required for cytokinesis from polyploid “titan” growth stages and yet retains an active site sequence signature. We propose that that a nuclease-independent function for Dis3L2 arose in an ancestral hyphae-forming fungus. This second function has been conserved across hundreds of millions of years, while the RNase activity was lost repeatedly in independent lineages.

## Introduction

Protein function evolves such that some descendants of an enzyme become “pseudoenzymes” with conserved structure but no catalytic activity (Murphy, Farhan, and Eyers 2017; Ribeiro et al. 2019). Distinct families of RNase enzymes regulate gene expression by catalytically degrading RNA (Houseley and Tollervey 2009), as part of a wider set of RNA-binding proteins (RBPs) that regulate all stages of the mRNA life cycle (Singh et al. 2015). Some functional RNA-binding proteins are pseudonucleases, in which inactivation of the nuclease active site was accompanied by, or preceeded by, gain of function in other domains. Pseudonucleases in animals include EXD1 (Yang et al. 2016), SMG5 (Glavan et al. 2006), *Maelstrom* (Chen et al. 2015), and *Exuperantia* (Lazzaretti et al. 2016). How could such changes in function have evolved? One possibility is that, first, the ability of a nuclease to bind RNA substrates was enhanced in other domains, as a secondary “moonlighting” function. Subsequently, the ancestral enzymatic activity was lost while the moonlighting activity was retained (Jeffery 2019). Understanding this order of events can help identify conserved activities underlying pleiotropic phenotypes.

### RNase-II family exoribonucleases

Members of the RNase II / Dis3 family of 3′-5′ exoribonucleases play important roles across the tree of life, including the founding member of the family, *E. coli* RNase II, and the essential Dis3/Rrp44 nuclease component of the eukaryotic RNA exosome (Dos Santos et al. 2018). Dis3L2 is a relative of Dis3 that specifically degrades poly(U)-tailed mRNAs, such as products of the terminal-U-transferases (Malecki et al. 2013), in *Schizosaccharomyces pombe*, a role conserved in mammalian Dis3L2 (Ustianenko et al. 2013). Dis3-family nucleases consist of two N-terminal cold-shock / OB-fold domains (CSDs), a central funnel-shaped domain that we refer to as RNII (also called RNB), and a C-terminal S1/K-homology domain. The nuclease activity is conferred by a magnesium ion at the centre of the RNII domain’s “funnel”. Four conserved aspartic acid (D) residues form a motif, DxxxxxDxDD (using single amino acid code, where x is any residue), that is conserved in all known active RNase II-family nucleases. The first, third and fourth D (equivalent to D201, D209 and D210 in *E. coli* RNAse II) are thought to be required for coordinating the magnesium ion (Zuo et al. 2006), while the second D hydrogen bonds to the 3′OH of the terminal base in the active site (Frazão et al. 2006). Experimental mutation of these conserved aspartic acids abolished the nuclease activity of RNII domains, including *E. coli* RNase II (Frazão et al. 2006; Zuo et al. 2006), *S. cerevisiae* Dis3/Rrp44 (Dziembowski et al. 2007; Schneider, Anderson, and Tollervey 2007), Human Dis3 (Tomecki et al. 2010), Human Dis3L1 (Staals et al. 2010; Tomecki et al. 2010), *Arabidopsis thaliana* Dis3/Rrp44 (Kumakura et al. 2016), and *S. pombe* Dis3L2 (Malecki et al. 2013). Thus, any RNase II homolog lacking some or all of these catalytic residues is likely to lack the conventional nuclease activity and may be assumed to be a pseudonuclease.

### The ascomycete Ssd1/Sts5 family of inactive RNase II-like proteins

Ascomycete yeasts contain additional conserved RNase II-like pseudonucleases: Ssd1 (*S.cerevisiae*; *Sc*Ssd1) and Sts5 (*S.pombe*; *Sp*Sts5) lack the conserved catalytic residues of the RNII domain and act as RNA-binding proteins that repress translation (Uesono, Toh-e, and Kikuchi 1997; Jansen et al. 2009). *Sc*Ssd1 was discovered due to synthetic lethality in combination with cell cycle mutants (Sutton, Immanuel, and Arndt 1991; Wilson et al. 1991). Deletion or truncation of *Sc*Ssd1 has pleiotropic effects, including reduced tolerance of stresses arising from ethanol (Avrahami-Moyal, Braun, and Engelberg 2012), heat (Mir, Fiedler, and Cashikar 2009), calcium (Tsuchiya et al. 1996), the kinase inhibitor caffeine (Parsons et al. 2004), and multiple chemicals that stress the cell wall (Kaeberlein and Guarente 2002; Mir, Fiedler, and Cashikar 2009; López-García et al. 2010). Ssd1 homologs are required for virulence in ascomycete fungal pathogens of humans and plants, including *Aspergillus fumigatus* (Thammahong et al. 2019), *Candida albicans* (Gank et al. 2008), *Colletotrichum lagenarium* and *Magnaporthe grisea* (Tanaka et al. 2007). Finally, *Sc*Ssd1 was recently shown to be required to support the survival of aneuploid yeast, although the mechanism remains unclear (Hose et al. 2020). Since full-length *Sc*Ssd1 binds RNA without detectable degradation (Uesono, Toh-e, and Kikuchi 1997), these pleiotropic effects of Ssd1 loss presumably reflect the loss of RNA-binding, rather than nuclease activity.

*Sc*Ssd1 and *Sp*Sts5 were reported to act as translational repressors of specific mRNAs involved in cell growth and cytokinesis (Jansen et al. 2009; Nuñez et al. 2016). Moreover, a conserved motif was identified in the RNAs targeted by *Sc*Ssd1 and *Sp*Sts5 (Hogan et al. 2008; Nuñez et al. 2016), strongly indicating that the RNA-binding surface is also highly conserved. *Sc*Ssd1-mediated mRNA repression connects to the Regulation of Ace2 and Morphogenesis (RAM) network via the NDR-family protein kinase Cbk1 (Du and Novick 2002; Jorgensen et al. 2002): Ssd1 deletion suppresses the lethality of Cbk1 deletion. *Sc*Ssd1 is phosphorylated by Cbk1 at its N-terminus (Jansen et al. 2009), and Cbk1 regulation is conserved to *C. albicans* Ssd1 (Lee et al. 2015). Similarly, *S. pombe* Sts5 is regulated by the Cbk1 homolog Orb6, and deletion of the RNA-binding protein suppresses defects arising from an inactive kinase (Nuñez et al. 2016).

Here we ask, how are pseudonucleases such as Ssd1 and Sts5 related to Dis3-family enzymes, and when did the common ancestor to Ssd1 and Sts5 lose its nuclease activity? Our phylogenetic analysis establishes that Ssd1 is the least diverged homolog of Dis3L2 in Saccharomycete yeasts, despite its lack of an active site. We show that the active site was lost on at least four separate occasions in fungi, while the cold-shock domains are highly conserved across both active and inactive homologs, in most branches of dikarya and mucoromycota. We predicted that the non-nuclease function of Ssd1 is conserved beyond ascomycota, and verified this by demonstrating a requirement for Ssd1 in cytokinesis in polyploid “titan” but not euploid yeast of the basidiomycete yeast *Cryptococcus neoformans*.

## Results and Discussion

### Ascomycete RNase-II-family pseudonucleases descend from Dis3L2

To understand the evolution of *Sc*Ssd1 and *Sp*Sts5, we first checked pre-computed databases of protein homology. The PANTHER protein homology database includes *Sc*Ssd1 and *Sp*Sts5 within a single Dis3L2 phylogeny (PTHR23355:SF9; (Mi et al. 2010)). The most parsimonious interpretation is that modern Ssd1 and Dis3L2 proteins are the descendants of a single eukaryotic ancestor. The OrthoDB hierarchical homology database clusters *Sc*Ssd1 with Dis3L2 and Dis3 proteins in both eukarya and and fungi (groups 1104619at2759 and 67258at4751; (Kriventseva et al. 2019)). The OrthoDB group containing *Sc*Ssd1 and *Sp*Sts5 in ascomycota (group 109571at4890) also includes a group of homologs in ascomycete filamentous fungi, such as *Histoplasma capsulatum*, with an active site sequence signature. However, the active site has been lost in all the least diverged homologs in the saccharomycotina (OrthoDB group 8134at4891). This implies that the ancestral ascomycete (~650MYA (Lücking et al. 2009)) had an active Dis3L2-like RNase and that the active site was lost in its descendant in the ancestral saccharomycete (~500 MYA (Prieto and Wedin 2013)).

### Reconstructing RNase II families in opisthokonts and amoebozoa

To map Dis3L2 evolution beyond fungi, we next performed a BLASTP search (Sayers et al. 2020) against *Sc*Ssd1, *Sp*Dis3L2, and *Hs*Dis3L2 from 76 phylogenetically representative species. We focused on representative fungi with sequenced genomes including major model organisms, edibles, and pathogens, along with some animals/metazoa, other holozoa and holomycota (Torruella et al. 2015). We included amoebozoa as an outgroup. We filtered the list of BLASTP homologs to have E-value 1 or less, and alignment length 200aa or more, and removed truncated sequences. We then aligned the curated full-length sequences with MAFFT (Katoh and Standley 2013), trimmed gaps at gap threshold 0.1 with trimAl (Capella-Gutiérrez, Silla-Martínez, and Gabaldón 2009), created a Bayesian maximum likelihood tree using IQ-TREE 2 (Minh et al. 2020) running on the CIPRES science gateway (Miller et al. 2015), and plotted the tree using ggtree (Yu 2020), using ggplot2 (Wickham 2016) and tidyverse packages (Wickham et al. 2019) in R markdown (Xie, Allaire, and Grolemund 2018). Full data and code for these analyses are available (*doi:10.5281/zenodo.3950856*).

The maximum likelihood tree shows clear clusters for Dis3, Dis3L1, Dis3L2, mitochondrial homolog Dss1, and a branch of amoebozoan RNII-Like proteins (aRNIIL) that we do not pursue further (Figure 1A). This reproduces previous results on clustering of Dis3/Dis3L1/Dis3L2 homologs (Ustianenko et al. 2013), and is consistent with the reported domain structures of these proteins. For example, all Dis3 homologs have a N-terminal PIN endonuclease domain with conserved catalytic residues, and Dis3L1 homologs have a PIN domain lacking essential catalytic resides, as reported in human Dis3L1 (Staals et al. 2010; Tomecki et al. 2010). We note that Dis3 and Dis3L1 are each mostly single-copy, and Dis3L1 is found only in metazoa in both this analysis and in the PANTHER database (PTHR23355:SF35/SF30; (Mi et al. 2010)). Dss1 is absent from metazoa.

**Figure 1:**
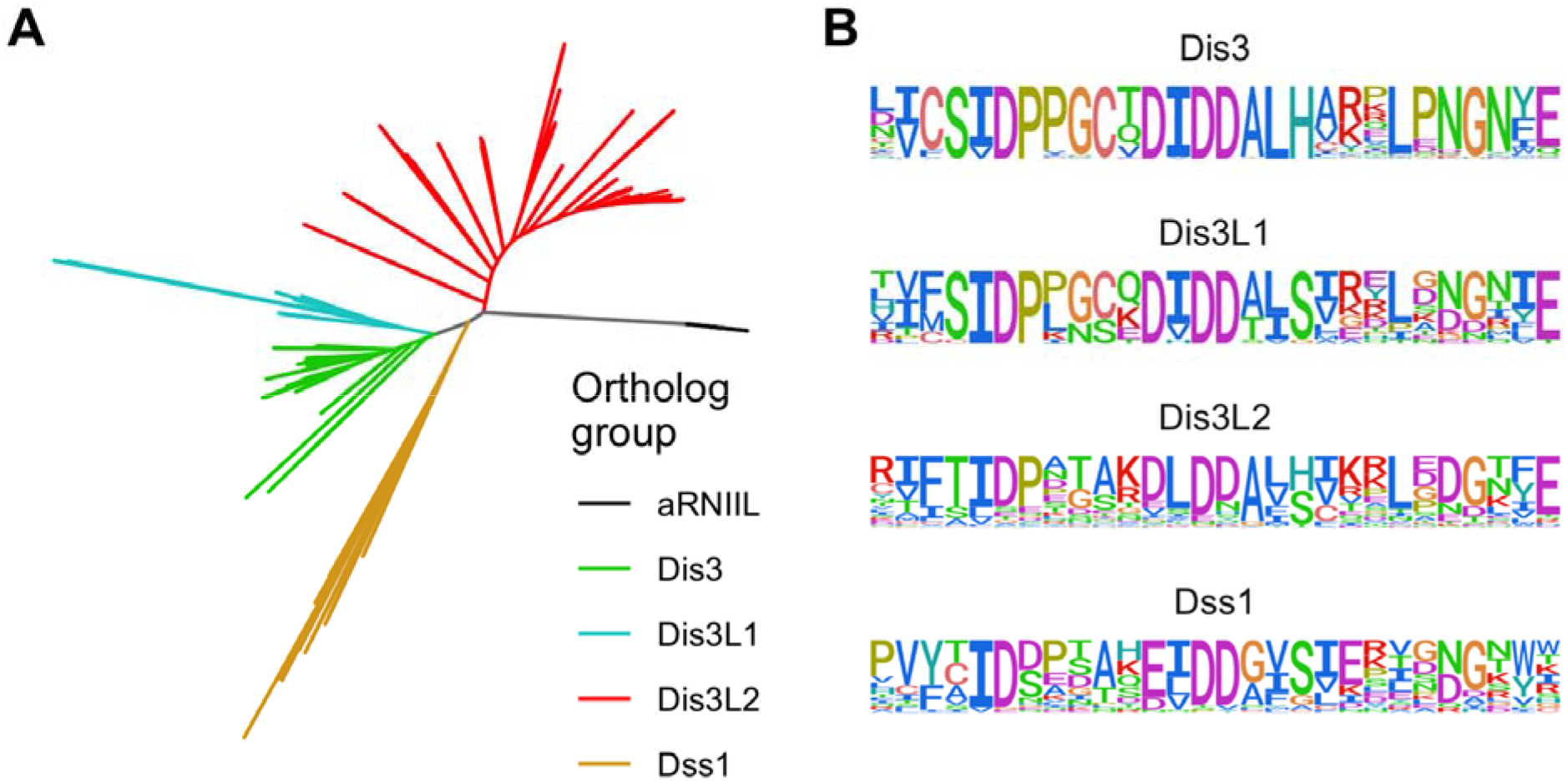
Phylogeny and active-site residues for Dis3 family enzymes in opisthokonta and amoebozoa. A, Phylogenetic tree of Dis3L2 and Ssd1 BLASTp homologs from 76 selected eukaryotes. Subfamilies are indicated in distinct colours: Dis3, Dis3L1, Dis3L2, Dss1, and amoebozoan RNII-Like proteins (aRNIIL). B, Consensus sequences (amino acid probability) for the RNII active site in Dis3, Dis3L1, Dis3L2, and Dss1 alignments.

### Related RNase II families have conserved nuclease active site signatures

We next computed consensus amino acid sequences for the active site of the larger clusters (Figure 1B), using the ggseqlogo package. This revealed a distinct active site signature for each subfamily. There is perfect conservation of the magnesium co-ordinating aspartic acids (D) in Dis3 (**D**PPgCx**D**I**DD**, where essential catalytic residues are bold, capital letters highly conserved, lower case letters indicate commonly occurring and x indicates any residue) and in Dis3L1 (**D**Pxxxx**D**I**DD**). However, both signatures for Dis3L2 (**D**Pxxxx**D**L**DD**) and Dss1 indicate that alternative residues appear in the conserved positions within this dataset. This indicates that both Dis3L2 and Dss1 lineages include some family members that are probable pseudonucleases, beyond *Sc*Ssd1. Furthermore, Dss1 shows a highly conserved E residue within the active site signature (**D**xxxxx**E**L**DD**), indicating that this conservative change can be tolerated in some active RNase II nucleases.

We sketch key features of the Dis3L2/Ssd1 family, including the position of the active site, in Figure 2A.

**Figure 2:**
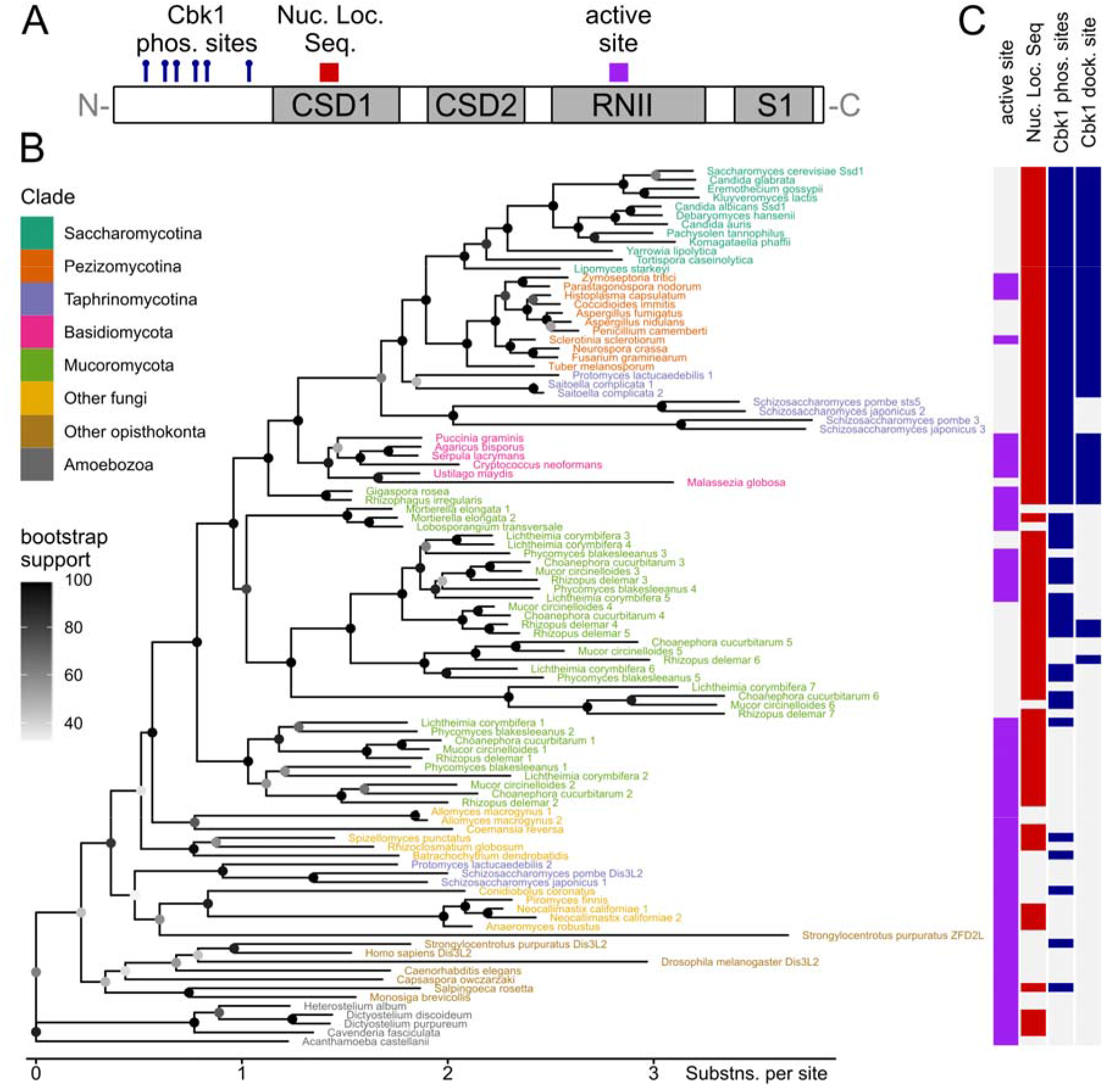
Evolution of the Dis3L2/Ssd1 family in fungi and relatives. A, Schematic of features found in Dis3L2 and Ssd1 family proteins. B, Phylogenetic tree of Dis3L2 family proteins, excluding N-termini aligned to *Sc*Ssd1 residues 1-337. Proteins are labeled by the species name coloured by clade, with a further identifier where there are multiple paralogs. Note that homologs from taphrinomycotina are in widely separated groups, e.g. S. pombe Dis3L2 and S. pombe Sts5. C, Features of Dis3L2/Ssd1 family proteins shown aligned with their position in the phylogenetic tree in B. For example, all homologs in Saccharomycotina have no active site, a nuclear localisation sequence, Cbk1 phosphorylation sites and a Cbk1 docking site. See text for details; full information with sequences, sequence identifiers, feature calculation and feature counts, is in the supplemental information.

### The Dis3L2 tree largely matches fungal species phylogenies

To examine the evolution of Dis3L2 homologs more closely, we generated a new multiple sequence alignment on the Dis3L2 cluster identified above, using the more accurate local pair option in MAFFT (Katoh and Standley 2013). We then computed the tree as previously, after removing the poorly-aligned N-terminus corresponding to *Sc*Ssd1 1-337, that is predicted to be unstructured (Figure 2B). As expected, the phylogeny of Dis3L2 homologs mostly follows species-level phylogenies as assembled from multiple genes (Ren et al. 2016). Most species in metazoa, saccharomycotina, pezizomycotina, basidiomycota, and chytridiomycota, have one homolog each, while mucoromycota have multiple homologs, reflecting repeated whole-genome duplications in this clade (Corrochano et al. 2016). We did not find any Dis3L2 homologs in microsporidia or cryptomycota, which are early-diverged fungi with reduced genomes and an intracellular parasitic lifestyle (James et al. 2013). Surprisingly, homologs in taphrinomycota are placed in two widely separated groups: *Sp*Dis3L2 seems to have diverged slowly with respect to basal opisthokonts, while *Sp*Sts5 clusters with other ascomycete homologs, but with a longer branch length that indicates faster sequence divergence. Repeated analyses with different gene lists and alignment parameters confirmed this wide separation (data not shown), although the exact placing of the *Sp*Dis3L2 group is poorly resolved, as indicated by the low bootstrap values.

To shed light on the evolution of Ssd1/Dis3L2 function, we next computed features of the (untrimmed) aligned protein sequences and displayed them alongside homologs in the tree (Figure 2C). An “active site signature” is identified where the three magnesium-co-ordinating Ds are in place in the RNII domain (Figure 2A). A classical nuclear localization signal was previously characterized in a loop in CSD1 of *Sc*Ssd1 (Kurischko et al. 2011); equivalently placed conserved sequences are identified. We identified regulatory Cbk1 kinase phosphorylation sites in the N-terminal region from the consensus Hxxxx[ST], including at least one positive amino acid (K or R) in the central xx residues, and the Cbk1 phosphorylation-enhancing docking site from its consensus sequence [YF]x[FP] (Gógl et al. 2015). The distribution of these features is not uniform across the Dis3L2 family (Figure 2C).

### The Dis3L2 active site signature is lost in at least four independent fungal lineages

All Dis3L2 homologs examined from amoebozoa, metazoa, and early-diverging chytridiomycota have the active site signature, indicating that the ancestral Dis3L2 was a nuclease (Figure 2C). The distribution of active site signatures on the phylogenetic tree indicates at least four independent losses of the active site in fungal Dis3L2s. First, the entire budding yeast Saccharomycotina clade has inactive Ssd1/Dis3L2 homologs, indicating a loss of the active site in an ancestor of the entire clade. Second, filamentous fungi in the pezizomycotina have a mix of active- and inactive-signature homologs, indicating a loss of the active site in the ancestor of *Aspergillus* and potentially also independently in the ancestor of *Tuber melanosporum*. Third, the dandruff-causing dermatophyte *Malassezia globosa* has an inactive homolog, despite clustering within the active homologs of its basidiomycete relatives. The active site is also lost in all sequenced members of genus *Malassezia* (data not shown). Fourth, in some groups of post-genome-duplication mucoromycota homologs the active site has been lost, e.g. *Rhizopus delemar* 5/6/7, despite closely related homologs with an intact active site signature, e.g. *Rhizopus delemar* 3. Indeed, our phylogenetic tree shows with high confidence that the active site has been lost on multiple branches diverging from the extant active-signature *Rhizopus delemar* 3.

Most Dis3L2 homologs contain a positively-charged nuclear localisation sequence in a loop in CSD1, similar to *Sc*Ssd1, suggesting that nuclear localisation is common in this family regardless of nuclease activity. One exception is *Sp*Dis3L2 and its active homologs in taphrinomycotina, which have lost the NLS signature in this location.

### Regulation of Dis3L2 by kinases is conserved beyond dikarya

Phosphorylation sites and docking sites recognised by the cell wall biogenesis kinase Cbk1 in *Sc*Ssd1 are conserved in almost all dikarya and many mucoromycota Dis3L2 homologs (Figure 2C). Cbk1 phosphorylation sites are a paradigmatic example of short linear motifs that are conserved in otherwise fast-diverging disordered regions (Zarin et al. 2019). Indeed, in the poorly aligned N-terminal domain of Dis3L2, multiple Hxxxx[ST] phosphorylation motifs stand out as strikingly conserved, e.g. 7 motif instances in *S. cerevisiae*, 8 instances in *C. neoformans*. A partial exception are members of the *Sp*Sts5 group that have 2 phosphorylation motifs but lack the Cbk1 docking site, and which are regulated by the diverged Orb6 kinase (Nuñez et al. 2016). It was previously noted that Cbk1 sites are conserved in saccharomycotina and pezizomycotina (Jansen et al. 2009), and this analysis argues for even deeper conservation of Dis3L2/Ssd1 regulation.

Ssd1/Dis3L2 regulation could also involve further post-transcriptional modifications. *S. cerevisiae* Ssd1 phosphorylation *in vivo* was reported to require the cyclin dependent kinase Cdk1 (Holt et al. 2009; Albuquerque et al. 2008). However, this requirement is likely to be indirect because Cdk1 regulates Cbk1 through a signaling cascade (Mancini Lombardi et al. 2013). Measurements in cell lysates failed to detect Ssd1 as a direct Cdk1 target (Ubersax et al. 2003). We did not pursue Cdk1 regulation further here because the two Cdk1 consensus sites [S/T]Px[K/R] on *Sc*Ssd1 are not conserved in our alignment, and the Cdk1-dependent sites indirectly identified *in vivo* overlap with verified and conserved Cbk1 phosphorylation sites.

### Ssd1 cold-shock domains are highly conserved in dikarya and mucoromycota

We next examined the domain conservation patterns of Dis3L2/Ssd1 domains (Figure 3). We computed pairwise percent amino acid identity in the trimmed MAFFT alignments for CSDs 1 & 2 (*Sc*Ssd1 338-659) and the RNII domain (*Sc*Ssd1 689-1014), shown in Figure 3 in the same sequence order as the tree in Figure 2B. The CSDs are highly conserved within saccharomycota, pezizomycotina, and basidiomycota, and quite highly conserved between these clades. CSDs are much less well conserved in basal fungi, metazoa and amoebozoa, contrasting with the higher conservation of RNII domains in these Dis3L2 nucleases. By contrast, the RNII domains are well-conserved within active-signature nucleases in the basidiomycota, with the exception of pseudonucleases in *Malassezia*.

**Figure 3:**
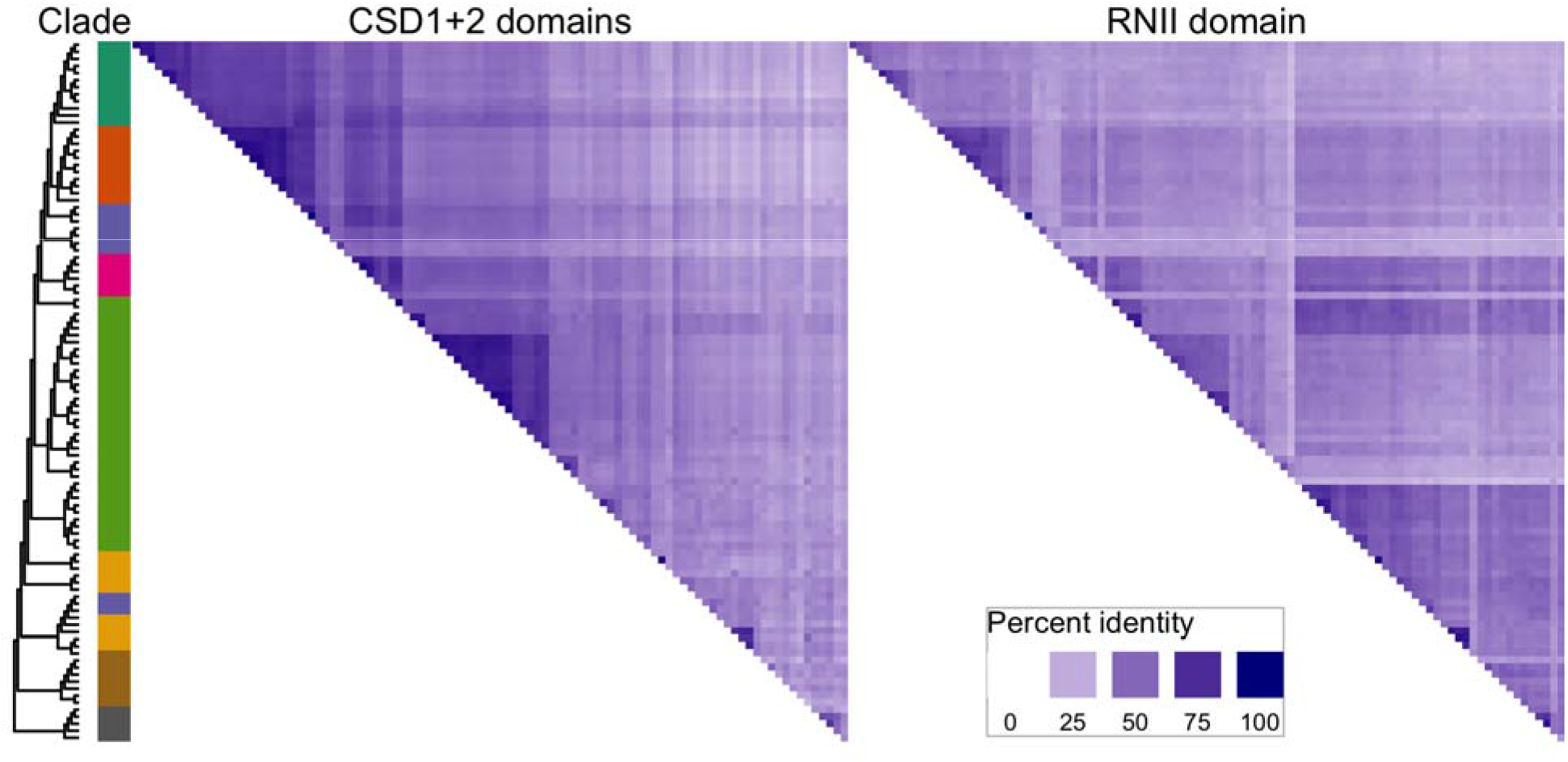
Conservation of Dis3L2-family domains in fungi and relatives. Heatmap shows percent identity of alignments within specific domains CSD1 and CSD2 considered together, and RNII domain, with darker blues indicating higher conservation. For example, dark blue patches at top left of CSD1+2 indicate that these domains are highly conserved within Saccharomycotina, compared to the lighter colours in the corresponding region for RNII indicating lower conservation. Cladogram and clade colouring is repeated from figure 2, as these are the same sequences in the same order.

These results may explain why previous reports focusing on nuclease activity in the RNII domain have argued that *S. cerevisiae* lacks a Dis3L2 homolog (Malecki et al. 2013; Lubas et al. 2013), as the active site region of the RNII domain is particularly diverged. By contrast, phylogenetic analysis and conservation of the CSDs place Ssd1 unambiguously as the least-diverged homolog of Dis3L2 in Saccharomycotina.

### Ssd1 has a conserved role in cytokinesis in the basidiomycete yeast *Cryptococcus neoformans*

To examine conservation of function associated with conservation of features between ascomycota and basidiomycota, we analysed the Ssd1/Dis3L2 homolog in the basidiomycete yeast *C. neoformans*. *Cn*Ssd1 is of interest because it retains a nuclease active site signature but also has features related to inactive Ssd1 homologs (nuclear localization signal, Cbk1 docking and phosphorylation sites). We used the *ssd1*Δ/CNAG_03345 ORF deletion from the Madhani laboratory deletion collection in the H99 background (Chun and Madhani 2010). Previous analysis of *ssd1*Δ found a slight growth defect, but no impact on yeast-phase morphogenesis (Gerik et al. 2005); we were able to replicate these findings during yeast phase growth in rich medium (data not shown). *C. neoformans* display two different morphologies: a yeast-phase budding phenotype and a much larger, polyploid “titan” morphology (>10 μm), that is associated with aneuploidy and virulence (Zaragoza and Nielsen 2013; Zhou and Ballou 2018). *In vitro* titan induction of wild type cells (*SSD1*) yields a mixed population of both yeast-phase and titan cells (Figure 4A) (Dambuza et al. 2018). Under this condition, the *ssd1*Δ strain shows defects in cytokinesis specifically in titan cells (Figure 4A). Among *ssd1*Δ cells, the average cell diameter was roughly 2 μm greater than *SSD1* cells, and the majority of mother cells >10μm had 2 or more daughters associated with the bud neck (p<0.0001, Mann-Whitney U test; Figure 4B). We observed no morphological or growth defects in yeast-phase cells that are also present during titan induction (Figure 4A).

**Figure 4:**
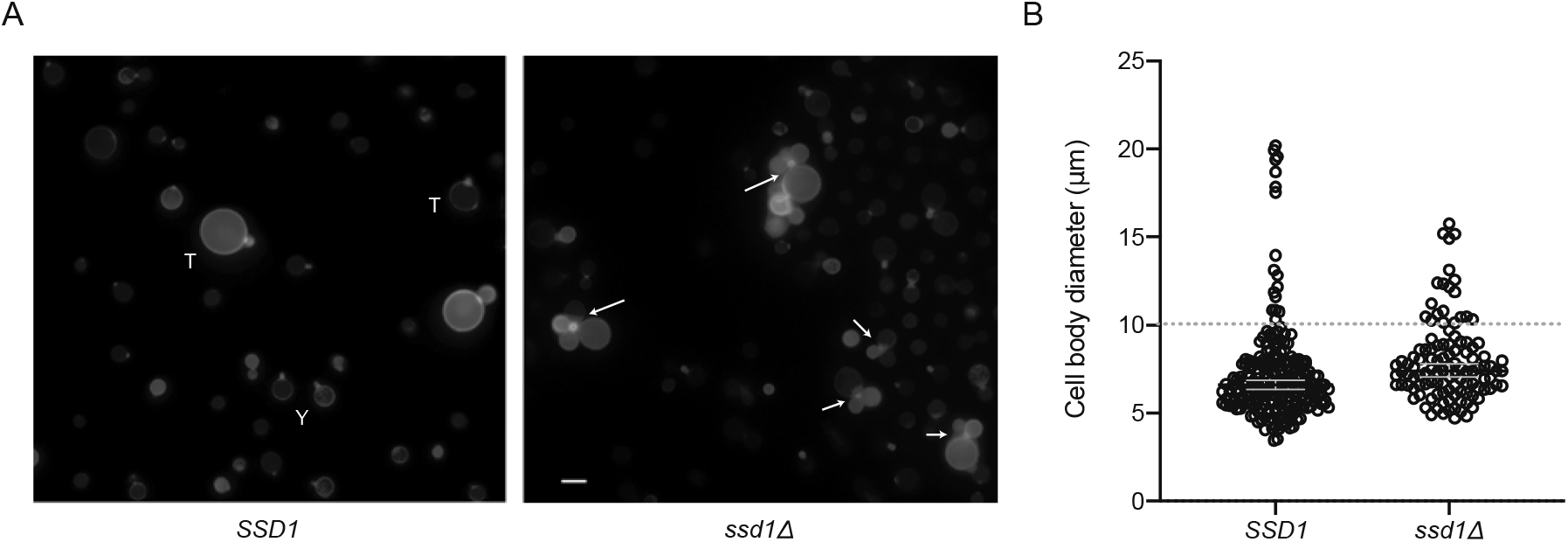
*Cryptococcus neoformans* Ssd1 is required for cytokinesis from polyploid titan-phase growth but not yeast-phase growth. SSD1 (wild-type strain H99) and ssd1Δ *C. neoformans* were grown in titan-inducing conditions as previously described (Dambuza et al. 2018). A, cells were stained for chitin using 0.1 μg/ml calcofluor white and imaged using a Zeiss AxioImager at 63x. Scale bar indicates 10 μm. Y indicates representative yeast cells, T indicates representative titan cells, and arrows indicate cells with abnormal cytokinesis. Among WT mother cells, none were observed with more than one bud; among ssd1Δ cells, the majority of mother cells >9μm had 2 or more daughters associated with the bud neck. B, The diameter of >100 cells was measured and analysed by Mann-Whitney U test for non-parametric data (p<0.0001). Median diameter and 95% CI are shown. All cells in 5 randomly selected frames were measured. Data are representative of three independent repeats but only a single experimental repeat is shown.

Our observation suggests a conserved role of *Cn*Ssd1 in cytokinesis, but that is either specialised to polyploid titan morphology, or that is redundant with other regulators during yeast-phase growth. These findings are consistent with those of Hose et al. showing that loss of ScSsd1 function is lethal for aneuploid cells but not euploid cells (Hose et al. 2020). Overall, the data suggest that these conserved functions are not related to nuclease activity, but may instead be connected to Cbk1 regulation. The Cbk1 kinase is required for cytokinesis from yeast-phase growth in *C. neoformans* (Walton, Heitman, and Idnurm 2006), and we speculate that this reflects Cbk1-mediated regulation of RNA binding by Ssd1.

### Evolution of an inactive RNA-binding protein from an ancestral nuclease via a bifunctional intermediate

Our work suggests a scenario where an ancestral Dis3L2 nuclease evolved a second RNA-binding function in a common ancestor of dikarya and mucoromycota. This ancestral fungus was likely developing a multicellular lifestyle involving spatially extended hyphal growth (Kiss et al. 2019). Given the reported role of modern-day Ssd1 homologs in mRNA localisation and translational control, we speculate that this role was played by Ssd1 in the ancestral hyphal fungus.

Nucleases can display weak RNA binding activity on surfaces distal to the active site, as a means of increasing their affinity for substrates. These additional sites can adapted during evolution, leading to bifunctionality in these enzymes, which can be followed by loss of the nuclease activity. Our results show multiple independent losses of nuclease activity in fungal homologs of Dis3L2, subsequent to the emergence of a conserved function of the cold-shock domains. The opisthokont exosome is a more extreme example, where 6 core PH nuclease-like proteins have all lost activity compared with the archaeal exosome and bacterial PNPase (Houseley and Tollervey 2009). These core proteins are pseudonucleases with a role in RNA binding. Nuclease activity in the opisthokont exosome is now restricted to the Dis3/Rrp44 subunit, or to the homologous Dis3L1 subunit of the metazoan cytoplasmic exosome (Staals et al. 2010; Tomecki et al. 2010). Even there, the PIN domain of Dis3 is an active endonuclease yet the PIN domain of Dis3L1 lacks nuclease activity while ensuring Dis3L1 binds to the core exosome as a “pseudonuclease domain”. Thus, pseudonucleases are a common feature of complexes that bind and regulate RNA.

Although the evidence is unambiguous that RNII domains lacking catalytic D residues are inactive, proteins that retain these residues are not necessarily active, as access to the active site could be blocked by other means. For example, active site residues might be retained in inactive enzymes, where access to the active site is blocked by mutations that occlude the RNA binding channel. The strong conservation of the active site and RNII domain in some clades, such as most basidiomycota, argues that retained nuclease activity is likely in these clades. Future experiments will have to address if diverged Dis3L2 homologs are active nucleases *in vivo* or *in vitro*, and if the nuclease activity is required for wild-type cell growth.

Lastly, we note that the canonical nuclease function of Dis3L2s requires its canonical substrates: RNAs that have poly(U) tails added by terminal U-transferases such as *S. pombe* cid1 and cid16, and human TUT1, TUT4, TUT7 (Yashiro and Tomita 2018). In the absence of terminal U-transferase activity, there would be few poly(U)-tailed substrates, removing selective pressure to retain Dis3L2’s terminal U-targeted nuclease activity. In this context, a bifunctional RNase/RBP would be unconstrained to evolve into a monofunctional RNA-binding pseudonuclease. Conversely, if terminal U-targeted nuclease activity were lost, there might be pressure against retaining an active TUTase, to avoid accumulation of poly(U)-tailed substrates. Supporting the coevolution of Dis3L2 and TUTase enzymes, TUTases homologous to *Sp*cid1/cid16 are present in fungal clades with active-signature Dis3L2 such as taphrinomycotina, most basidiomycota, mucoromycota, and chytridiomycota (PANTHER:PTHR12271:SF40; OrthoDB:264968at4751), but absent from prominent clades lacking active Dis3L2, such as most saccharomycotina.

Overall, our analysis identifies extant fungal Ssd1 homologs as descendants of the Dis3L2 family of 3′-5′ exoribonucleases, identifies the CSDs as highly conserved features across dikarya that are likely to perform conserved functions related to aneuploidy and cytokinesis, and raises new questions about the interaction of these domains with client RNAs.

## Acknowledgments

We thank Gemma Atkinson for essential and generous advice on phylogenetic methods. We thank David Tollervey, Marah Jnied, Laura Tuck, and members of the Wallace lab for discussions and comments on the manuscript. E.R.B and E.W.J.W. are each supported by Sir Henry Dale Fellowships jointly funded by the Wellcome Trust and the Royal Society [211241/Z/18/Z to E.R.B., 208779/Z/17/Z to E.W.J.W.]. A.G.C is a Wellcome Senior Research Fellow [200898] in the Wellcome Centre for Cell Biology [203149]. We thank Hiten Madhani for making the *C. neoformans* gene deletion collection available, funded by NIH grant (R01AI100272).

Data and code underlying the evolutionary analysis is available at *doi:10.5281/zenodo.3950856*.

